# Isolation of a novel heterodimeric PSII complex via strep-tagged PsbO

**DOI:** 10.1101/2022.06.15.496312

**Authors:** Jan Lambertz, Jakob Meier-Credo, Svetlana Kucher, Enrica Bordignon, Julian D. Langer, Marc M. Nowaczyk

## Abstract

The multi-subunit membrane protein complex Photosystem II (PSII) catalyzes the light-driven oxidation of water and with this the initial step of photosynthetic electron transport in plants, algae, and cyanobacteria. Its biogenesis is coordinated by a network of auxiliary proteins that facilitate the stepwise assembly of individual subunits and cofactors, forming various intermediate complexes until fully functional mature PSII is present at the end of the process. In the current study, we purified PSII complexes from a mutant line of the thermophilic cyanobacterium *Thermosynechococcus vestitus* BP-1 in which the extrinsic subunit PsbO, characteristic for active PSII, was fused with an N-terminal Twin-Strep-tag. Three distinct PSII complexes were separated by ion-exchange chromatography after the initial affinity purification. Two complexes differ in their oligomeric state (monomeric and dimeric) but share the typical subunit composition of mature PSII. They are characterized by the very high oxygen-evolving activity of approx. 6,000 µmol O_2_· (mg Chl·h)^-1^. Analysis of the third (heterodimeric) PSII complex revealed lower oxygen-evolving activity of approx. 3,000 µmol O_2_· (mg Chl·h)^-1^ and manganese content of 2.7 (± 0.2) per reaction center compared to 3.7 (± 0.2) of fully active PSII. Mass spectrometry and time-resolved fluorescence spectroscopy further indicated that PsbO is partially replaced by Psb27 in this PSII fraction, thus implying a role in the repair of the complex.

## 1 Introduction

Cyanobacterial photosystem II (PSII) is a large membrane protein complex characteristic of oxygenic photosynthesis. It catalyzes the light-driven oxidation of water and the reduction of plastoquinone, thus injecting electrons into the photosynthetic electron transport chain and releasing dioxygen as a byproduct. PSII is present as monomeric and dimeric complexes, with each monomer consisting of at least 20 protein subunits and many different cofactors like chlorophylls, quinones, carotenoids, lipids, and the unique Mn_4_CaO_5_ cluster, which is responsible for water oxidation [1,2]. Together with other membrane protein complexes like photosystem I (PSI), the cytochrome-b_6_f complex (cyt-b_6_f), and ATP synthase, photosynthetic organisms produce ATP and NADPH, which are used to generate the primary biomass on our planet.

The main redox cofactors of PSII are coordinated by the central D1 and D2 subunits [3], whereas light energy is additionally collected by the antenna proteins CP43 and CP47 and transferred to the reaction center (RC) chlorophylls P_680_ [4]. Upon charge separation at the RC, an electron is transferred via pheophytin to a bound plastoquinone molecule (Q_A_), and then passed to the mobile electron carrier Q_B_. The resulting electron-hole in P_680_^+^ is filled by an electron from an adjacent tyrosine residue (Tyr), which in turn oxidizes the oxygen-evolving complex (OEC) [5]. The catalytic cycle of the OEC consists of 5 different redox states (S_0_-S_4_), where the index indicates the number of stored oxidation equivalents required to split two water molecules [6]. The OEC is shielded from the lumen by the extrinsic subunits PsbO, PsbU and PsbV, three soluble proteins characteristic of active cyanobacterial photosystem II. Although not essential for PSII activity, oxygen evolution and stability are drastically reduced when they are not present [7].

PSII biogenesis and repair involve a spatially separated, stepwise assembly process facilitated by numerous auxiliary protein factors, which are not part of the active protein complex [8–10]. However, the characterization of PSII assembly intermediates is hampered by their low abundance and intrinsic instability. Several proteins have been reported to be involved in PSII assembly, including the well-characterized auxiliary factors Psb27 and Psb28 [11–14]. Although neither is essential for PSII maturation, both play a role in the process. Psb27 facilitates OEC formation at the PSII donor side [11,15], while Psb28 protects PSII from photodamage by structurally modifying the PSII acceptor side during assembly [11,16]. The D1 subunit is most susceptible to photodamage and is therefore frequently replaced by newly synthesized protein [17]. The exact nature and sequence of intermediate steps is not yet fully understood, but the process is likely initiated by the destabilization of the D1-DE loop and the loss of the bicarbonate [18]. This might be followed by the release of the extrinsic proteins and binding of Psb27 at the lumenal side of the inactive PSII dimer [19,20]. In the next steps, PSII is monomerized, the CP43 module is removed and the damaged D1 subunit is replaced by a newly synthesized copy [17]. Reassembly of the complex and photoactivation of the OEC appears to proceed as in biogenesis. Moreover, the structural changes induced by either PSII subunit or auxiliary protein binding, often affect the intrinsic redox properties of PSII, for example by modulating the Q_A_ redox potential, thus promoting safe recombination to protect the proteins during assembly and/or repair [11,21,22].

In the present study, we purified a novel, heterodimeric PSII intermediate via strep-tagged PsbO from the thermophilic cyanobacterium *Thermosynechococcus vestitus* BP-1 (formerly known as *Thermosynechococcus elongatus* BP1). The intermediate is potentially involved in PSII repair, as biochemical characterization, activity measurements, EPR analysis and time-resolved fluorescence kinetics revealed that one of the OECs is disrupted and PsbO is partially replaced by Psb27. In addition, two highly active PSII species (monomer and dimer) were isolated, which in principle are well suited for further structural and functional studies to gain a deeper understanding of this very unique enzyme.

## 2 Results

A *T. vestitus* mutant line was generated, in which the Twin-Strep-tag (TS-tag) sequence [23] was linked to the N-terminus of PsbO (Figure SI 1) to facilitate the isolation of active PSII by Strep-tag affinity chromatography (AC). Three different PSII complexes were separated by subsequent ion-exchange chromatography (IEC) (Figure 1a). The first fraction contains only monomeric PSII, with a relative contribution of 23 ± 5.6% to the PSII total (Figure 1b, Table 1). The other two complexes are dimeric, with relative proportions of 69 ± 3.9% and 7.6 ± 1.7%, respectively. Oxygen evolution measurements revealed a high activity of the monomeric and the major dimeric fraction, with 5,154 ± 1,084 and 6,078 ± 512 µmol O_2_·mg Chl^-1^·h^-1^, respectively (Table 1, Figure 1c). Based on the oxygen evolution and oligomeric state, we assigned these fractions to active monomer (aM) and active dimer (aD). Strikingly, the less abundant dimeric fraction, which we termed semi-active dimer (saD), has an approximately 50% lower activity of 3,083 ± 711 µmol O_2_·mg Chl^-1^·h^-1^.

**Table 1:** Overview of PSII species that were isolated via TS-PsbO. Data of ≥ 3 independent preparations are shown.

|  | active Monomer | active Dimer | semi-active Dimer |
| --- | --- | --- | --- |
| Abbreviation | aM | aD | saD |
| Oligomeric state | monomeric | dimeric | dimeric |
| Relative amount (%) | $23 \pm 5.6$ | $69 \pm 3.9$ | $7.6 \pm 1.7$ |
| Oxygen evolution activity<br>($\mu\text{mol O}_2 \cdot (\text{mg Chl}^{-1} \cdot \text{h})^{-1}$) | $5,154 \pm 1,084$ | $6,078 \pm 512$ | $3,083 \pm 711$ |
| Mn/RC* | $3.38 \pm 0.15$ (91%) | $3.72 \pm 0.21$ (100%) | $2.69 \pm 0.24$ (72%) |

\*In parenthesis the percentage of Mn ions with respect to aD

**Figure 1:**
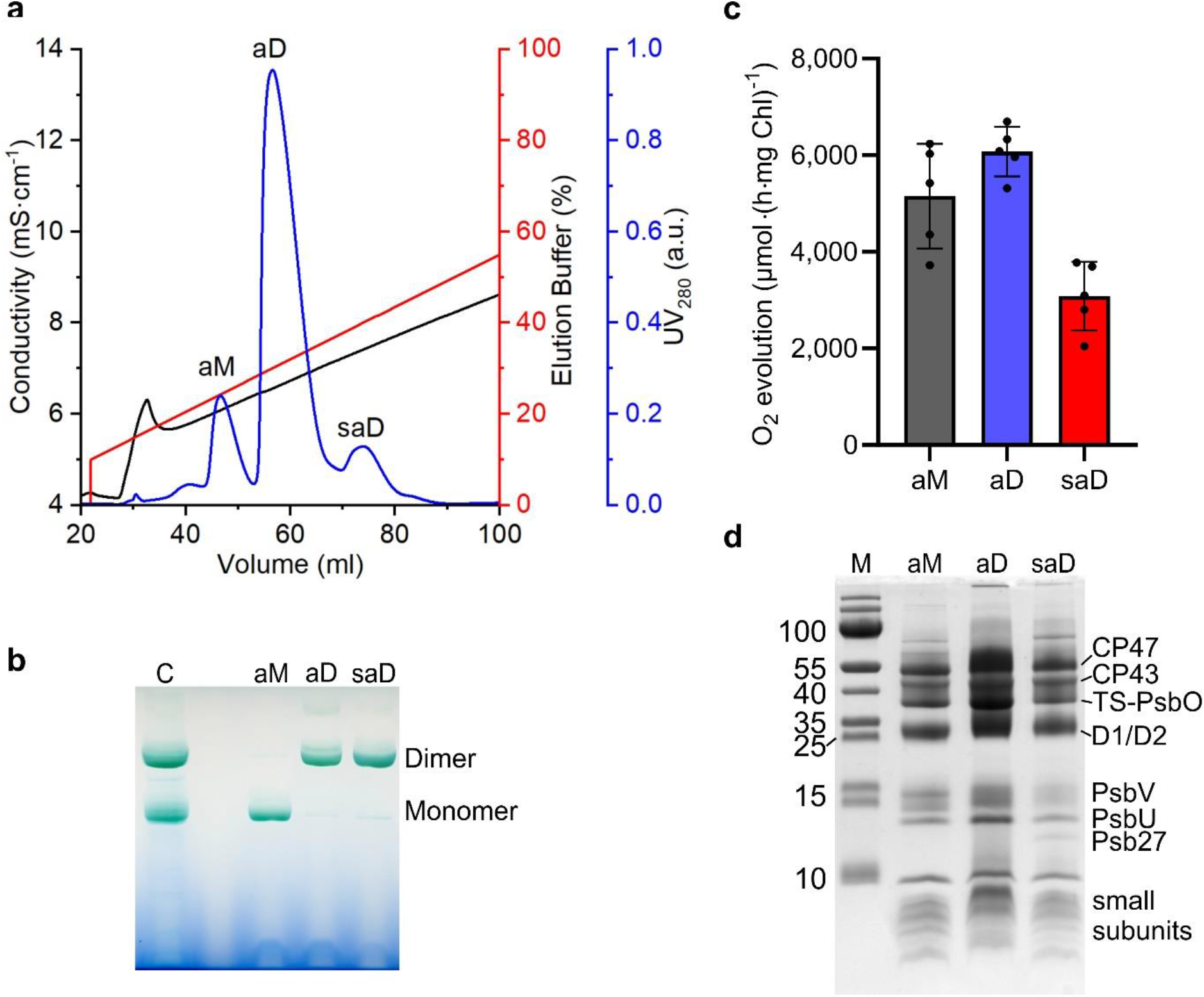
Isolation of three different, active PSII complexes. **a)** Ion-exchange chromatogram of the PSII preparation after Strep-tag AC. Absorbance A_280_ (blue), conductivity (black) and gradient of elution buffer (red) plotted against the volume. **b)** BN-PAGE of purified PSII complexes after IEC. C: PSII monomer and dimers isolated with a TS-tag on CP43 [24] **c)** Oxygen evolution rates of PSII complexes in presence of the electron acceptors DCBQ (1 mM) and ferricyanide (5 mM). Each data point represents the measurement of a separate PSII preparation, which was isolated from independent cell cultures (biological replicate) and the average of at least two technical replicates. **d)** SDS-PAGE of purified PSII complexes after IEC. M: Size reference (Prestained PageRuler #26616, Fisher Scientific)

The protein composition of the three PSII complexes was investigated by SDS-PAGE, LC-MS/MS and MALDI-TOF MS. The results revealed that stoichiometric amounts of all major subunits characteristic of active PSII are present in aM and aD (Figure 1d), including the extrinsic subunits PsbU and PsbV. The signal of PsbO shifted close to the CP43 band due to the additional mass of the TS-Tag (Figure 1d, Figure SI 2). Notably, in saD, an additional band at ∼ 13 kDa is present, which we assigned to the auxiliary factor Psb27 by LC-MS/MS (SI File 1), whereas PsbU and PsbV seem to be less abundant. This finding was unexpected, as Psb27 was previously described to bind exclusively to inactive monomeric and dimeric PSII [11,14,19,20]. However, Psb27 might be still an indicator for inactive PSII complexes, if one monomer in saD is inactive, binds Psb27 and lacks PsbO, PsbU as well as PsbV, whereas the other is fully active due to the presence of a functional Mn_4_CaO_5_ cluster and the extrinsic proteins. The recent cryo-EM structures of Psb27-containing PSII complexes revealed that some of the low molecular weight subunits (LMWP) are missing [11,20]. Therefore, we analyzed aM, aD and saD by LC-ESI-MS/MS and MALDI-TOF MS for the presence of LMWPs (Figure 2, Table 2, SI File 2, SI Table 1). The results confirm that most of the known LMWPs are present in all three complexes of our PSII preparation, except for Psb30 and PsbY, which seem to be lost during sample preparation.

**Table 2:**
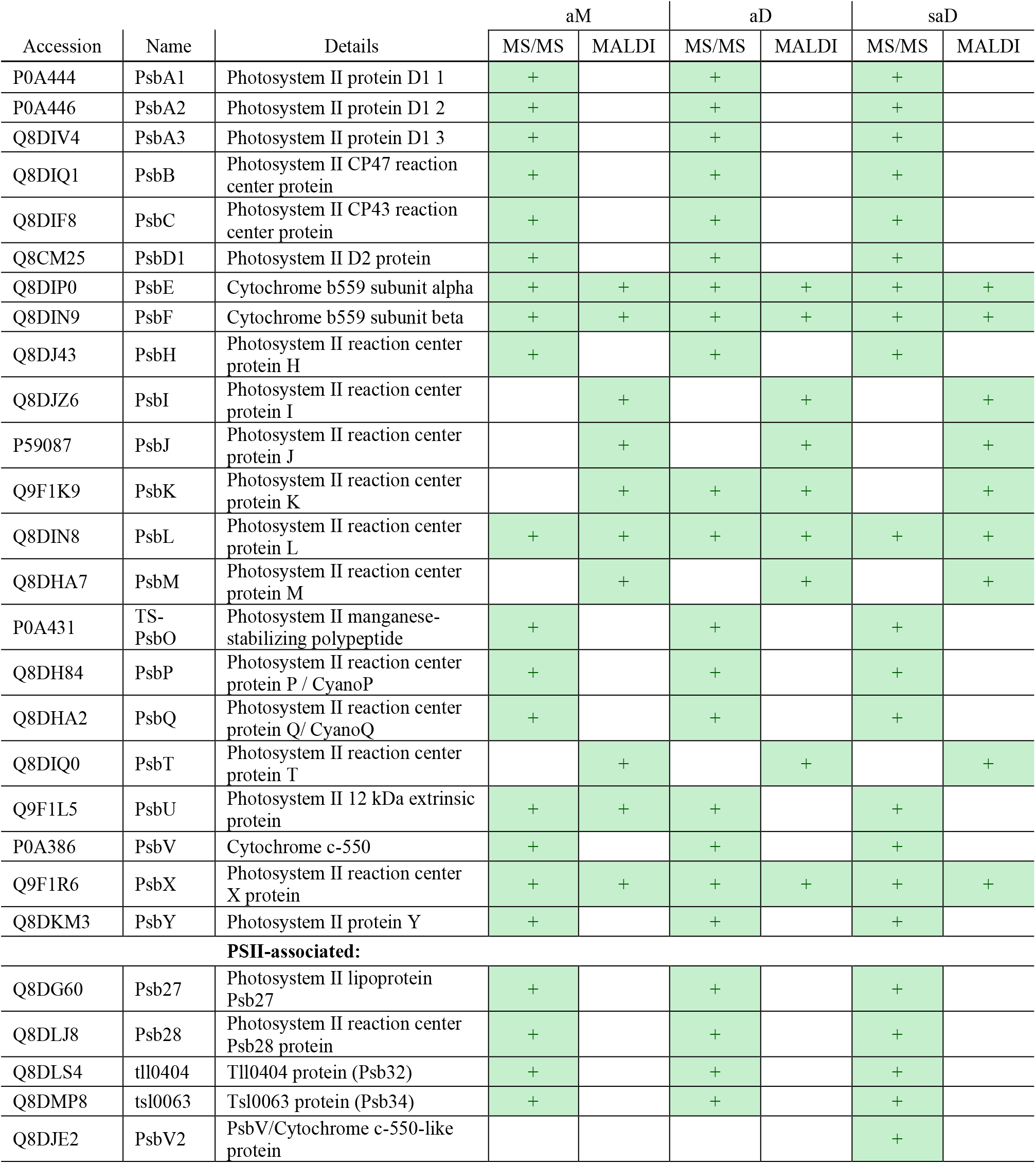
Proteins identified by LC-MS/MS and MALDI-ToF-MS. The subunits of PSII and PSII-associated proteins are shown. See SI Tables for a full list of identified proteins.

**Figure 2:**
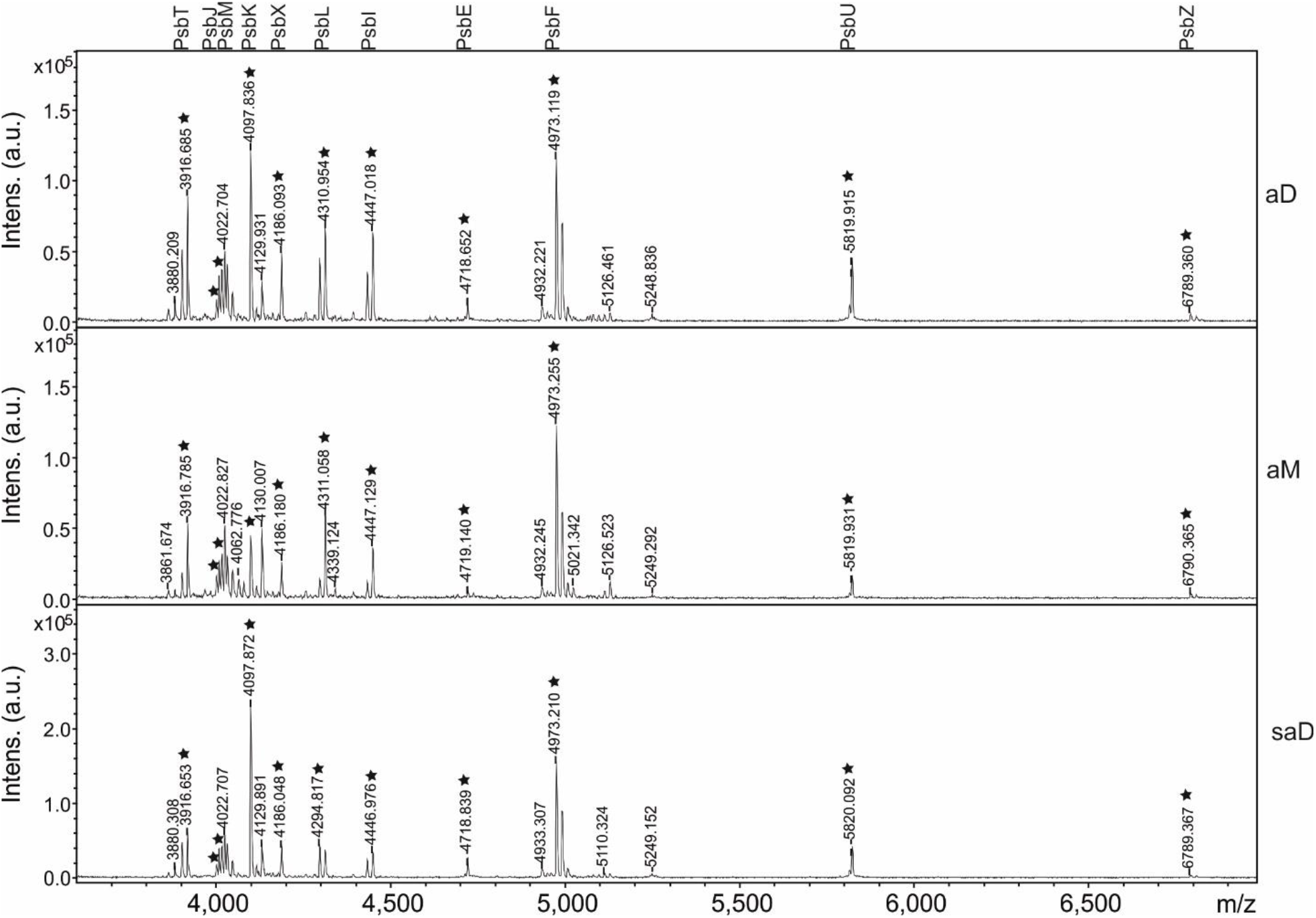
MALDI-TOF spectra of the small proteins of PSII. Representative graphs of 4 technical replicates for each complex are shown. a) m/z range from 3,600 to 7,000; Annotation of the peaks was done according to [25,26]. Stars indicate modified peptides (see also SI Table 1).

LC-MS/MS analysis further revealed proteins related to PSII in all complexes (e.g., Psb28, Psb32, Psb34, etc.), which are not visible on SDS-PAGE or identified by MALDI-ToF MS. It is likely that they are only present in substoichiometric amounts (Table 2, SI MS Table 2). Interestingly, PsbV2 (*tll1284*) was only found in saD. These proteins might belong to low abundant intermediate PSII complexes that were co-purified via PsbO-TS.

The presence of Psb27 is usually indicative of reduced manganese content in the corresponding PSII complexes [11,20,27], although it has been shown that a dimeric complex involved in PSII repair retains about four manganese atoms per reaction center (RC) in an inactive conformation if Psb27 is bound [19] and a recent study describes a Psb27-PSII complex with a fully assembled OEC [28]. We analyzed the Mn-content of the three PSII complexes by electron paramagnetic resonance (EPR) spectroscopy and confirmed the presence of almost 4 Mn atoms (3.72 ± 0.21) per RC in aD (Figure 3, Table 1, Figure SI 4-6, SI Table 2-3). The Mn-content in aM is slightly reduced (3.38 ± 0.15), probably due to damage of PSII monomers during preparation or due to a fraction of still not activated PSII, whereas saD contains about 72% (2.69 ± 0.24) of the usual manganese content per RC of fully active PSII, which corresponds to 5.75 vs 8 Mn ions in aD. As four Mn atoms are required for each reaction center to be operational [2], one of the two Mn clusters of saD must be fully assembled to explain the water-splitting activity of saD. Consequently, the other monomer contains only 1-2 Mn-atom, probably bound to the high-affinity binding site, which binds the first Mn-atom of the cluster [11]. The presence of more than one Mn is possibly due to a fraction of dimers containing more Mn-ions in one of the RC. In conclusion, the determined Mn-content of saD explains its oxygen-evolving activity, which is about half of aD on a per-chlorophyll basis, thus supporting a heterodimeric structure of saD.

**Figure 3:**
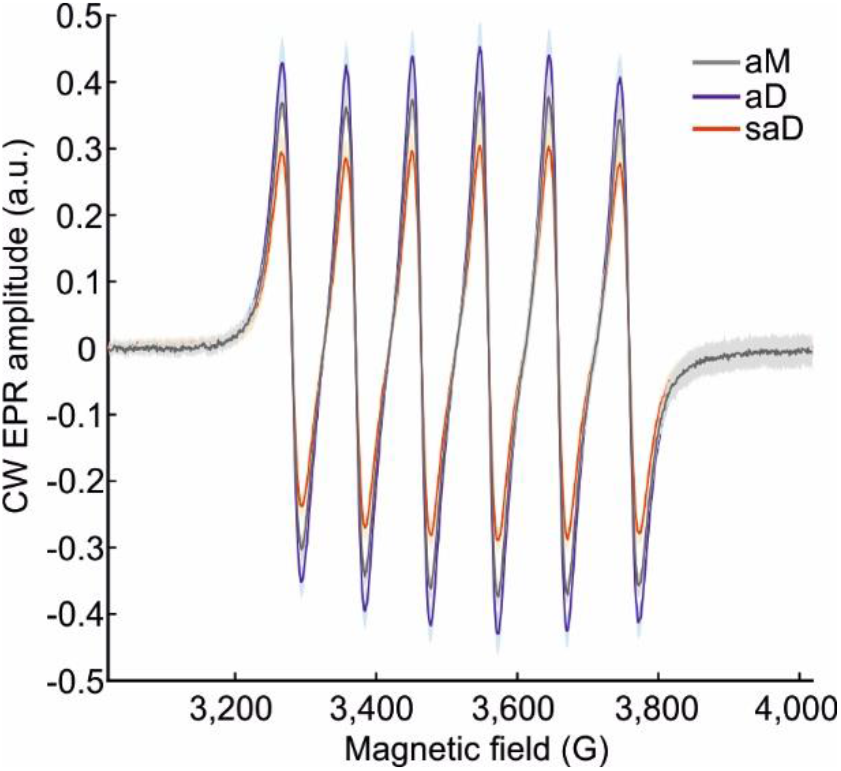
Manganese quantification. **a)** Offset and background-corrected X-band (9.86GHz) continuous wave (CW) EPR spectra of supernatants of aM, aD and saD (C_RC_=17.6 µM) after acid extraction [27,29]. The solid lines represent an average spectrum of several biological and technical repeats of raw data shown in SI Figures 5 and 6 with standard deviations shown as shaded areas.

The photochemical properties of the complexes were further investigated by time-resolved flash-induced Chl a fluorescence decay measurements. A single short, saturating light flash is used to induce charge separation at the reaction center chlorophylls and subsequent reduction of Q_A_ in most of the complexes. The fluorescence signal decays in a multistep process and variations in the kinetics indicate differences in the internal electron transfer chain. As shown in Figure 4a, all three complexes exhibit the typical three-phase decay [11,30]. However, deconvolution of the data (equation 1) for each complex revealed differences in amplitudes and half times of the different reactions (Table 3).

**Table 3:** Chl a fluorescence decay kinetics. Relative Amplitudes (Amp (%)) of the total fluorescence signal and the corresponding half-times (t_1/2_) are shown. Differences to 100% are due to rounding. Values in brackets indicate the quality of the fit (for details, see methods).

|  | fast phase |  | middle phase |  | slow phase |  |
| --- | --- | --- | --- | --- | --- | --- |
| | Amp (%) | $t_{1/2}$ ($\mu\text{s}$) | Amp (%) | $t_{1/2}$ (ms) | Amp (%) | $t_{1/2}$ (s) |
| no treatment |  |  |  |  |  |  |
| aM | 44 | 678 ($\pm 47$) | 13 | 37.8 ($\pm 7.2$) | 44 | 12.5 ($\pm 0.7$) |
| aD | 55 | 324 ($\pm 41$) | 29 | 2.8 ($\pm 0.5$) | 16 | 1.6 ($\pm 0.2$) |
| saD | 44 | 887 ($\pm 58$) | 19 | 51 ($\pm 6.0$) | 36 | 7.6 ($\pm 0.5$) |
| DCBQ |  |  |  |  |  |  |
| | Amp (%) | $t_{1/2}$ ($\mu\text{s}$) | Amp (%) | $t_{1/2}$ (ms) | Amp (%) | $t_{1/2}$ (ms) |
| aM | 41 | 182 ($\pm 21$) | 29 | 2.25 ($\pm 0.3$) | 30 | 80.7 ($\pm 6.6$) |
| aD | 54 | 213 ($\pm 16$) | 24 | 3.39 ($\pm 0.4$) | 22 | 125.5 ($\pm 12$) |
| saD | 43 | 408 ($\pm 149$) | 28 | 8.00 ($\pm 3.9$) | 29 | 183.5 ($\pm 56$) |
| DCMU |  |  |  |  |  |  |
| | Amp (%) | $t_{1/2}$ (ms) | Amp (%) | $t_{1/2}$ (s) | Amp (%) | $t_{1/2}$ (s) |
| aM | 10 | 460 ($\pm 84$) | 40 | 5.3 ($\pm 0.4$) | 50 | 102 ($\pm 12$) |
| aD | 1.2 | 13.8 ($\pm 22$) | 60 | 3.8 ($\pm 0.1$) | 39 | 142 ($\pm 21$) |
| saD | 14 | 37.5 ($\pm 5.8$) | 39 | 2.6 ($\pm 0.2$) | 46 | 101 ($\pm 12$) |

**Figure 4:**
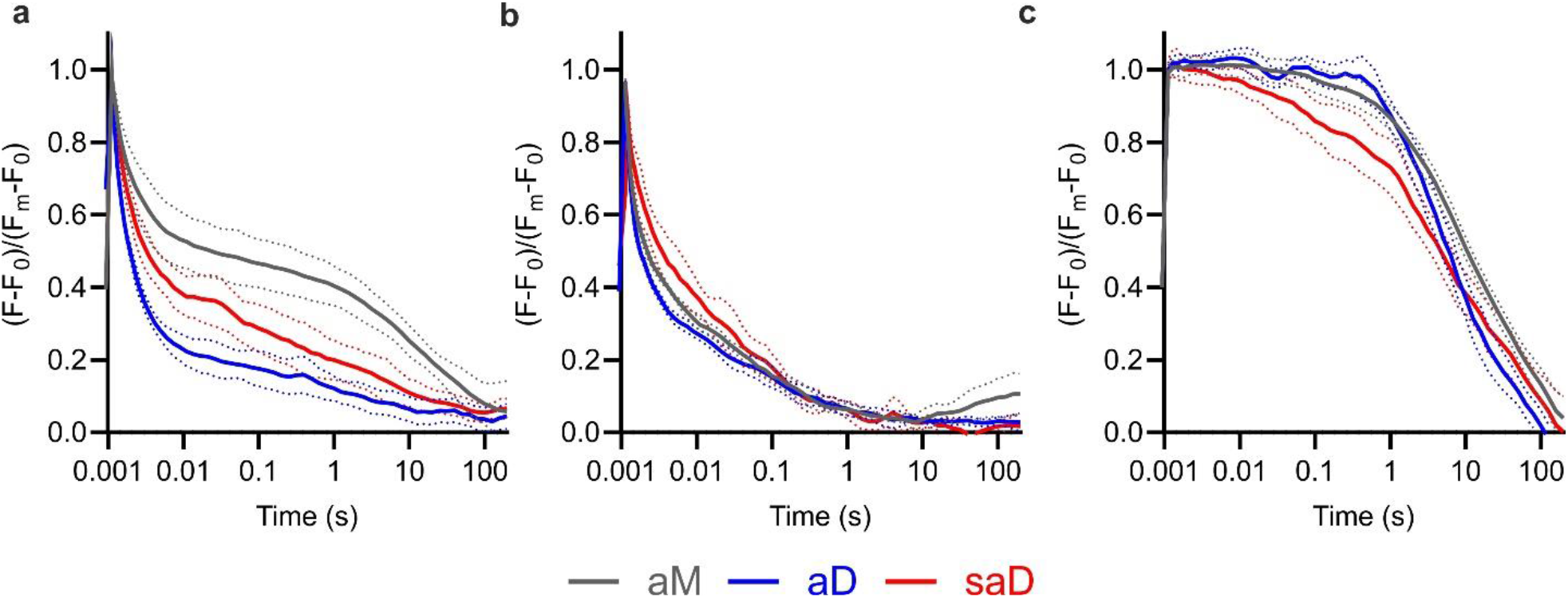
Chl a fluorescence decay measurements of isolated PSII complexes. The Chl a fluorescence decay of isolated PSII complexes after a single, saturating light flash was measured without treatment **(a)** and in presence of 20 µM DCBQ **(b)** or 20 µM DCMU **(c)**. Data were normalized to the signal before the light flash (F_0_) and after the light flash (F_m_). Curves represent averages of at least three replicates, dotted lines indicate standard deviation.

The fast decay phase originates from Q_B_-reduction [30] and 55% of the complexes in aD and 44% of the complexes in aM and saD (Figure 4a, Table 3) contribute to this fast forward electron transfer. The calculated half-life time of the fast phase of aD (t_(1/2)_= 324 µs) is comparable with values determined for whole cells [18,30]. The decay is nearly three times slower (t_(1/2)_ = 887 µs) in saD and two times slower in aM (t_(1/2)_= 678 µs), either indicating an impaired forward electron transfer due to the lack of Q_B_ or increased charge recombination in these complexes. A similar deceleration can be observed for the middle phase (Figure 4a), which represents the interaction with improperly bound Q_B_ [31] or, in the case of intact membranes, the interaction with the PQ-pool [30]. In aD, 29% of the photosystems contribute to this slower reduction of Q_B_ with a half time of 2.8 ms, comparable to intact *Synechocystis* cells [30]. Together with the fast phase, 84% of all aD complexes facilitate forward electron transfer. In contrast, the middle phase has an amplitude of only 13 or 19% and is slowed down to 37.8 or 51 ms in aM or saD, respectively (Table 3).

Due to the lower Mn content of saD, half of the complexes in the heterodimer are likely incapable of oxygen evolution and lack a functional Mn cluster, which promotes safe recombination by an increased Q_A_ redox potential [32]. Thus, P_680^+^_Q_A^-^_recombination in the 0.8-1 ms time scale [30,33], and Tyr_Z^+^_Q_A^-^_ recombination in the middle phase (t_(1/2)_ = 80-120 ms) [33,34] may further contribute to the decay kinetics in saD (and to a lesser extent in aM), thereby leading to a slower fluorescence decay.

To examine if the slower kinetics originate from impaired forward electron transfer due to less co-purified Q_B_, fluorescence decay kinetics were measured in presence of the artificial electron acceptor DCBQ. The addition of DCBQ (20 µM final concentration) accelerated the fast phase of both aM and aD to ∼182 and 213 µs, respectively, and for aM the amplitude of the middle phase was increased to 29% in the expanse of less recombination in the slow phase (Table 3 and Figure 4b). The half-life time of the middle phase was drastically accelerated in aM to a value of 2.25 ms, which is comparable to that of aD. The minimal changes in the fluorescence decay kinetics reveal a high Q_B_-content in aD, whereas it seems lower in aM. In saD, both the fast and middle phases are accelerated by DCBQ addition but are still slower compared to aM and aD. The fluorescence decay kinetics are both decreased for the fast phase (t_½_ = 400 µs) and the middle phase (t_(1/2)_ = 8 ms), which indicates increased charge recombination in complexes incapable of Q_B_-reduction. The value determined for the middle phase seems too fast for Tyr_z_^+^Q_A_^-^ recombination and too slow for P_680_^+^Q_A_^-^ recombination [11,30,33–36]. However, binding of auxiliary factors or loss of Q_B_, bicarbonate or the OEC may accelerate Tyr_z_^+^Q_A_^-^ recombination to the lower millisecond range of about 10 ms [11,30,37]. This is likely the case for the inactive half of saD and the observed half-life time of the fluorescence decay is comprised of a mixture of the active and inactive half of the complex.

The fluorescence decay of the third (slow) phase is dominated by S_2_Q_A_^-^ recombination with intact Mn-clusters [11,28,30,38]. In aD, the half time is 1.58 s (S_2_Q_A_^-^), and slow recombination occurs in 16% of all complexes, while 10 times slower values were obtained for aM (44%, 12.5 s). Slower decay rates above ∼10 seconds have been attributed to recombination of “trapped” Q_A_^-^ in complexes with a stable electron donor or side-path donors like Mn^2+^, Tyr_D_ or cyt-b559 [28,36,39]. In saD, a mixture of the signals is present (36%, 7.6 s), which can be explained by the above-mentioned heterodimeric structure of the complex. Notably, the kinetics of the third phase are shifted to 80 – 180 ms for all three complexes in the presence of DCBQ (Table 3). This indicates that in acceptor saturated complexes, recombination with the S2-state or alternative electron acceptors is absent, whereas Tyr_z_^+^Q_A_^-^ recombination still occurs.

Next, fluorescence decay measurements in the presence of the PSII inhibitor 3-(3,4-dichlorophenyl)-1,1-dimethylurea (DCMU) were performed (Figure 4c) to examine the recombination effects in more detail. DCMU inhibits electron transfer from Q_A_ to Q_B_ and diminishes Q_A_ re-oxidation by forward electron transfer. The corresponding fluorescence data were deconvoluted by using equation (1) with either an exponential and hyperbolic decay [11,30], or two exponential and one hyperbolic decay [30], with the latter approach resulting in better fits for the data presented here (Figure SI 7).

DCMU addition diminishes the fast decay in aD, as seen in the reduction of the fast phase-amplitude to 1.2% (Table 3). This inhibitory effect of DCMU is even greater than previously observed [11,30,40], indicating the high amount of fully intact complexes, while in aM and saD 10 and 14% of the complexes, respectively, retain a fast decay. Although, while the half times of saD and aD are increased to 37.5 and 12 ms, those of aM are ten times slower (460 ms). However, the small amplitudes may impact the fitting, resulting in the inaccuracy of the time constant. As mentioned above, these reactions resemble recombination in complexes with an impaired donor site [30,37] and recently, a similar phase in a PSII monomer from *Nicotiana tabacum* with a half time of 50 ms was assigned to Tyr_z_^+^Q_A_^-^ recombination in complexes with an incomplete Mn-cluster [28].

The remaining signal decays with one exponential and one hyperbolic decay (Table 3, Figure 4c). The exponential (middle) decay is accelerated in saD with a half time of 2.6 s, compared to aM and aD with half times of 5.3 and 3.8 s, respectively. Similar values were obtained for Synechocystis cells (t_(1/2)_ = 3.2 s), where Mn-depletion further accelerated the recombination (t_(1/2)_ = 1.6 s) [36], consistent with the reduced Mn-content of saD. The low-second-scale phase represents S_2_Q_A_^-^ recombination in complexes with an intact OEC and without an alternative, stable donor [30], but its origin in Mn-depleted PSII is yet unclear. The slow hyperbolic decay of all three complexes lies in the minute scale and reflects recombination in complexes in presence of alternative electron donors, in which Q_A_^-^ is “trapped” [11,28,36]. In complexes with an intact Mn-cluster, this involves slow S_2_Q_A_^-^ recombination and, in the fraction of complexes without an intact Mn-cluster, alternative pathways like bound Mn-atoms [39].

We further investigated the variable fluorescence induction kinetics of aM, aD and saD to get more insights into their intrinsic electron transfer properties. Under continuous illumination, the accumulation of Chl a fluorescence can be traced and typically, the first ten seconds of the measurement reveal four distinct features (O-J-I-P, Figure 5). Beginning with the initial F_0_ or O-feature, reduced Q_A_ accumulates (J-feature), followed by full reduction to Q_A_^-^Q_B_^2-^ at F_m_ (P-Peak) [41–44]. The intermediary phase (I feature) is missing in systems without an intact membrane and is supposed to reflect the influence of the electric potential across the thylakoid membrane [43], or a heterogeneous PSII population [42]. For a semi-quantitative comparison, the data was evaluated using a three-exponential decay equation (2) to represent the different transitions [43,45]. We additionally analyzed the OJIP-traces using a two-exponential decay function based on equation (2), if applicable (Table 4), as we expect no intermediate feature due to the lack of a membrane environment in our samples. The active dimer - representing the PSII fraction with the highest abundance in the cell - reveals typical FI curves of pre-illuminated or DCMU treated cells [42,45], with 91% of Q_A_- reduction during the O-J transition and a half time of 33.22 ms (Figure 5a, Table 4). The J-I and I-P transitions have amplitudes with 8% (t_(1/2)_ = 0.35 s) and 1% (t_(1/2)_ = 5.10 s). By using two instead of three exponents, deconvolution only minorly affects the O-J transition (91% with t_(1/2)_ = 33.42 ms) and the J-P transition can be deconvoluted as a single phase with 9% of the total amplitude with t_(1/2)_ = 434 ms, resulting in a better fit accuracy. This value is remarkably similar to the time required for QB reduction (200-400 ms) [5].

**Table 4:** Induced Chl a fluorescence kinetic best fit values. Relative Amplitudes (Amp (%)) of the total fluorescence signal and the calculated half-times (t_1/2_) are shown. In some cases, more than one reaction was assigned to a single phase. The K and I-feature are exclusive for saD, and J for aM and aD. Values in brackets indicate the goodness of the fit (for details, see methods).

|  | no treatment |  |  |  | DCMU |  |  |  |
| --- | --- | --- | --- | --- | --- | --- | --- | --- |
|  | O-J/K |  | J-P/K-I-P |  | O-J/K |  | J-P/K-I-P |  |
| | Amp (%) | $t_{1/2}$ (ms) | Amp (%) | $t_{1/2}$ (s) | Amp (%) | $t_{1/2}$ (ms) | Amp (%) | $t_{1/2}$ (s) |
| aM | 22 (± 0.3) | 5.34 (± 0.080) | 8 (± 0.1) | 0.66 (± 0.008) | 93 (± 0.1) | 11.88 (± 0.02) | 7 (± 0.1) | 0.14 (± 0.002) |
| | 70 (± 0.3) | 29.78 (± 0.001) | | | | | | |
| aD | 91 (± 0.1) | 33.42 (± 0.066) | 9 (± 0.1) | 0.43 (± 0.009) | 98 (± 0.1) | 11.42 (± 0.04) | 2 (± 0.1) | 0.19 (± 0.029) |
| saD | 50 (± 3.0) | 17.79 (± 0.117) | 21 (± 0.2) | 0.09 (± 0.002) | 78 (± 0.2) | 10.44 (± 0.04) | 10 (± 0.4) | 0.11 (± 0.007) |
| | | | 29 (± 0.1) | 0.93 (± 0.008) | | | 13 (± 0.5) | 0.53 (± 0.021) |
| | 65 (± 0.1) | 25.36 (± 0.144) | 35 (± 0.2) | 0.64 (± 0.006) | | | | |

**Figure 5:**
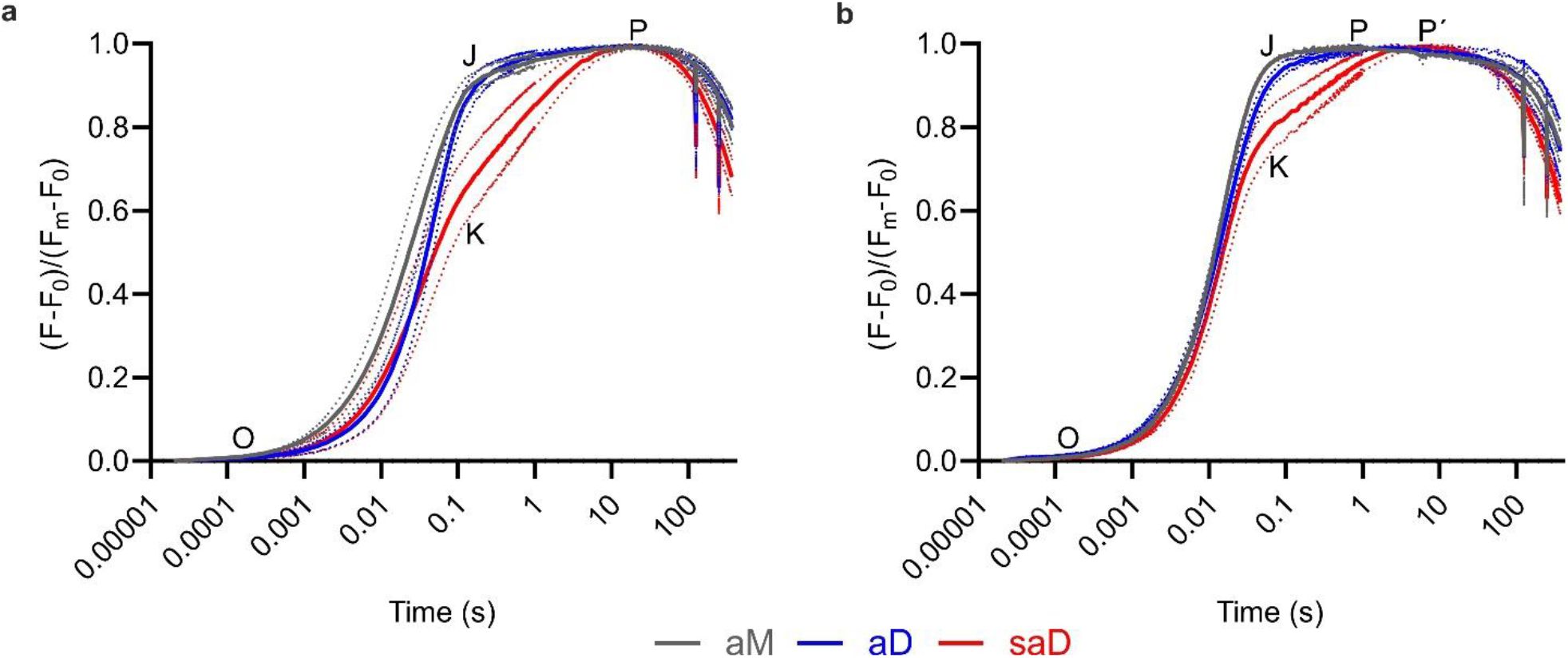
Variable fluorescence induction measurements of isolated PSII complexes. The fluorescence yield of the three complexes under continuous illumination in **(a)** absence and **(b)** presence of 20 µM DCMU reveal a lack of an I-, and the presence of a K-feature, normalized to the initial fluorescence F0 and Fm at the P-peak. The O, J, I, P and K features are labeled. P’ indicates F_m_ in saD. Curves represent averages of at least three replicates, dotted lines indicate standard deviation.

Consistent with the reduced Mn-content of saD, the kinetics (Figure 5a) exhibit the same features as described before for other PSII complexes with less manganese [21,46]. The induction kinetics of these complexes show a K-feature, which is characterized by a faster fluorescence rise in the 0.1 to 1 second time scale [44] due to a shift in the Q_A_ reduction equilibrium by an increased Q_A_ redox potential [21,46]. Best fits were obtained by separation of the K-I (21%, t_(1/2)_= 88.7 ms) and I-P (29%, t_(1/2)_= 0.926 s) transitions. Deconvolution of the data with a single reaction for the J-P transition (Table 4) resulted in the following values: 65% of the amplitude are defined by the O-J transition with t_(1/2)_= 25.36 ms, and 35% originate from the J-P-transition (t_(1/2)_= 642 ms), although the deconvolution of the latter is less accurate (Figure SI 7). Due to the heterodimeric character of saD, the K-feature is less pronounced than in artificially inhibited PSII or PSII-I [21]. The fluorescence rise of aM differs in the micro-and millisecond range (Figure 5a) compared to the other two complexes. In this case, a three exponential fit was used to deconvolute the data (Table 4, Figure SI 7). The half-life time of the first phase is very low (5.34 ms) and covers around 22% of the total amplitude. In combination with the “second” O-J phase (70% and 29.8 ms), the O-J transition contributes to 92% of the total amplitude with a combined t_(1/2)_ of 23.96 ms, which is ∼30% faster than in aD. The J-P transition has an amplitude of 8% and a half-time of 661 ms, which is about 52% slower than for aD.

To investigate whether the differences in the fluorescence induction kinetics were due to different amounts of Q_B_ in the samples, DCMU was added to a final concentration of 20 µM before the measurement (Table 4, Figure 5b). Because DCMU inhibits Q_B_ reduction, it mimics the lack of Q_B,_ which results in faster Q_A_^-^ accumulation. Best results were obtained for aM and aD with a two-exponential fit, whereas three reactions gave the best fitting result for saD (Figure SI 9). Inhibition of forward electron transfer leads to faster Q_A_^-^ accumulation with t_(1/2)_ = 11.88 and 11.42 ms for aM and aD, respectively. The differences in the O-J transitions, particularly the additional early increase in aM, are diminished. Thereby, the additional first phase in aM arises from differences in Q_B_-reduction, which can be explained by the partial lack of Q_B_ in aM resulting in an accelerated accumulation of Q_A_^-^ in a fraction of complexes even in absence of DCMU. Still, the contribution of the O-J transition in aM is reduced compared to aD (93% vs 98%). This indicates that in aD or aM, 2% or 7% of the complexes, respectively, have either impaired Q_A_-reduction or promoted recombination effects, which is consistent with the ∼9% reduced Manganese content of aM.

The kinetics of saD are less affected by DCMU-addition (Figure 5b) and, in contrast to aM and aD, best fits were obtained by using three exponentials (Figure SI 9). The O-K transition is increased by a factor of 1.25 to 78% of the total amplitude (t_(1/2)_ = 10.44 ms) and the K-I as well as I-P transitions contribute with 10% (t_(1/2)_ = 0.11 s) and 13% (t_(1/2)_ = 0.53 s) to the amplitude, respectively. Still, the proportion of the O-K transition is greater than observed for fully inactivated PSII [21], which is consistent with the heterodimeric structure of saD.

## 3 Discussion

We isolated three active PSII complexes from *T. vestitus* via TS-tagged PsbO by streptavidin affinity chromatography and subsequent ion-exchange chromatography. By targeting PsbO, nearly no inactive PSII complexes were purified, which are usually present in preparations with affinity tags fused to the core subunits [14,24,47]. The complexes are of high purity and all major subunits are present in stoichiometric amounts. The active dimer (aD) represents the major PSII fraction and catalyzes the reaction of approx. 52 O_2_·s^-1^ per RC (104 per PSII dimer), which indicates a turnover frequency of around 19 ms per dioxygen evolved. This is remarkably high compared to previous PSII preparations, especially in view of the high purity and homogeneity of the various complexes [14,24,36,48–52].

The Chl a fluorescence decay of aD revealed Q_B_ reduction rates remarkably similar to whole cells [30]. More than 80% of the complexes in aD maintained Q_B_ bound to the acceptor side during preparation, represented by the fast and middle phase. Moreover, they are barely affected by the addition of the alternative electron acceptor DCBQ. The fluorescence decay kinetics in presence of DCMU revealed remarkable integrity of the complex, as Tyr_Z^+^_Q_A^-^_ recombination indicative for a disrupted OEC was barely detected [11,30,34,37]. Typically, the Q_B_-reduction rates appear slower in the fast phase due to Tyr_Z_ recombination as a background reaction, either due to degradation or heterogeneity of the used material [11,30,34,37]. The slow phase decay in presence of DCMU also revealed fast S_2_Q_A_^-^ recombination in aD, which indicates that alternative electron donor pathways play a minor role in aD [11,28,39]. As shown in the following, the high stability and purity of the samples allow a precise assignment of different reactions also in the other two PSII fractions, which is usually limited due to the heterogeneity of signals and overlays of different reactions.

The monomeric PSII fraction (aM) contains less Q_B_, which drastically altered the fluorescence kinetics to a slower apparent Q_B_-reduction and a faster accumulation of Q_A_^-^. These characteristics were not observed in the presence of DCBQ or DCMU, resulting in signals comparable to those of aD. Based on our results, Q_B_-reduction as well as recombination processes are influenced by variations in Q_B_ content, which should be considered when comparing complexes.

The stabilization of Q_A_^-^ in aM, which manifests in slower S_2_Q_A_^-^ recombination [39], indicates a stronger influence of alternative electron donors in the monomeric PSII compared to the dimer. The vast majority of OECs in aM is intact, but the 15% lower oxygen evolution and the 9% lower Mn-content of aM compared to aD, as well as the presence of Tyr_z_^+^Q_A_^-^ recombination in ∼ 5% of the complexes indicate a subpopulation of PSII with a disrupted OEC. This might correspond to a nearly mature PsbO-PSII-complex without oxygen evolution, which is close to the photoactivation of the OEC. As Psb27 was also detected in substoichiometric amounts in aM by MS, a Psb27-PsbO-PSII intermediate would be feasible. The presence of PsbO in monomeric PSII isolated by tagged Psb27 [40], as well as new structural data [11,20] indicate that simultaneous binding of both proteins is possible. Further separation of the monomeric fraction might provide novel insights into this PSII subspecies. However, we cannot exclude the alternative explanation that PSII monomers with a disrupted OEC originate from lower stability, which results in degradation during the preparation procedure.

We propose that the semi-active dimer (saD) is a heterodimer consisting of an active and inactive half, as oxygen evolution requires a fully assembled Mn-cluster [2]. Consequently, one OEC must be disrupted (∼ 1-2 Mn) in saD, while the other needs to be intact (4 Mn) to explain the decreased Mn content and the reduced oxygen evolution (50%) compared to the active dimer (aD). This is supported by the Chl a fluorescence decay measurements, which revealed recombination of Q_A_^-^ with Tyr_Z+_ and the S2-state. Tyr_Z^+^_Q_A^-^_ recombination is a feature of complexes without an intact Mn-cluster, in which Tyr_Z_ cannot be re-reduced efficiently by transition into the S_2_-state [30,37]. As seen by the lack of Tyr_Z_ recombination in aD, this reaction is specific to saD, together with the clear S_2_Q_A_^-^ recombination, which is an exclusive feature of complexes with a functional OEC [30]. The slower S_2_Q_A_^-^ decay within ∼10 s represents recombination effects in presence of alternative electron donors like Mn-atoms or Tyr_D_, which prevent fast oxidation of Q_A_^-^ by recombination with the donor side [11,36,39]. The slower fast and middle phase of the fluorescence decay of saD can be indicative of a slower Q_B_ reduction instead of Tyr_Z^+^_Q_A^-^_ recombination, which has been observed in PsbO-or bicarbonate-free PSII [28,37]. Because of the reduced Mn-content and the recombination seen in presence of DCMU, the slower decay presumably represents a mixture of Q_B_ reduction and Tyr_z_^+^Q_A_^-^ recombination in the different moieties of the heterodimer.

Another indication of the heterogeneous nature of saD is the presence of a K-feature in the fluorescence induction data. This feature is indicative of Mn-depletion and - in contrast to the J-feature – it is only slightly affected by DCMU addition [21,46]. Compared to fully depleted PSII, the K-feature is less pronounced, and the O-K amplitude is increased by the addition of DCMU due to the presence of the active half. It would be expected that the O-K transition in saD can be deconvoluted into two reactions, but this was not possible, likely because the O-J/K transitions have very similar half times (Table 4). Remarkably, the transition to the P-peak revealed the presence of an I-inflexion in saD, but neither in aM nor aD. The inflexion may originate from the membrane potential [43], which can be excluded in the case of isolated complexes, or it originates from different PSII populations within the sample [42,53]. The lack of a distinct I-feature for aM and aD supports the former explanation, but its presence in saD suggests that the I-inflexion – at least partially – is a feature of PSII subpopulations, thereby of PSII heterogeneity in the cell or sample.

The heterodimeric structure of saD is further supported by the presence of Psb27, an auxiliary protein specific for complexes without the extrinsic subunits, often characterized by a lack of oxygen evolution and a reduced Mn-content [11,14,19,20,28,40]. Simultaneous binding of both Psb27 and PsbO to one RC has been proposed to be limited or short-lived [11,20,40], and because of the specificity of Psb27 for PSII intermediates, it presumably binds to the OEC-disrupted half of saD, while the extrinsic subunits maintain oxygen evolution in the other.

In contrast to PSII biogenesis, the sequential steps of PSII repair remain elusive. The proposed models assume six major steps [9,17,19,54,55]: i) Detachment of the extrinsic subunits and disassembly of the OEC, ii) Binding of Psb27, iii) Monomerization, iv) Detachment of CP43, v) Replacement of damaged subunits and vi) Re-assembly of the complex according to PSII biogenesis. In these models, saD might represent an early intermediate, in which one half is active and the other is repaired. Successive inactivation of both moieties would result in the formation of the inactive PSII dimer [19,20]. Alternatively, the inactive half of saD is close to photoactivation after repair or during PSII assembly. The imminent photoactivation would explain the low Mn content (1-2) compared to the inactive PSII dimer, which retains 3-4 Mn per RC in the presence of Psb27 [19]. In this case, detachment of Psb27 and binding of the extrinsic subunits would be the next steps besides manganese incorporation [10].

The exclusive detection of the PsbV-like protein PsbV2 in saD by mass spectrometry is an interesting finding. Photoautotrophic growth can be restored by overexpression of PsbV2 in PsbV deletion strains, but the strains retain their impaired growth under low Ca^2+^ and Cl^-^ concentrations, which are important for the functionality of the Mn-cluster [56]. In contrast, PsbV2 deletion strains revealed no distinct phenotype both under normal growth and calcium/chloride limiting conditions [57]. According to our data, PsbV2 might be involved in PSII repair and/or assembly, which is in agreement with its low abundance in the cell, typical for PSII assembly factors [58,59].

In summary, we isolated highly active monomeric and dimeric PSII complexes, which are suitable candidates for in-depth spectroscopic analysis due to their very low intrinsic heterogeneity and high stability. In addition, a novel, heterodimeric PSII complex was purified, which is presumably composed of an active half with all extrinsic subunits, and an inactive moiety with Psb27 and a partially assembled Mn-cluster. This semi-active dimer shares Chl a fluorescence characteristic of both active and inactive PSII complexes.

## 4 Experimental Procedures

### Generation and cultivation of *T. vestitus* strains

A Twin-Strep-Tag (SAWSHPQFEKGGGSSGGGSAWSHPQFEKGGGS) [23] was fused to the N-terminal region of PsbO, directly successive to K28 after the signal peptide sequence. The Plasmid with the modified PsbO DNA sequence including up- and downstream regions of 1,000 bp around the modified N-terminus was synthesized by Twist Bioscience (USA) and cloned into their high copy pTwist plasmid. A multiple cloning site (MCS) was introduced in the synthetic DNA-sequence 190 bp upstream of PsbO, in which a kanamycin resistance cassette was inserted by ligation (Figure SI 1). The plasmid was integrated into *T. vestitus* BP-1 wild-type by electroporation [60]. Segregation of the mutant line was done by gradually increasing the kanamycin concentration up to 150 µg/ml (Figure SI 1). *T. vestitus* cultures were cultivated at 45 °C in bg-11 medium in Snijder incubators or, for high volumes, in 25-L photobioreactors (Bioengineering) supplied with 5% CO_2_ [49]. Light intensity was increased depending on cell density from 50 to 300 µmol photons·m^-2^·s^-1^. Confirmation of the segregation was done by PCR of isolated genomic DNA using the primers GTACGCATTTGGACCTCATC upstream of the kanamycin cassette and GAGCAATGCGATAGGTTTGG in the PsbO-gene (Fig. SI 1). Cells from 25-L cultures were concentrated to 1 L using an Amicon® DC10 LA system, followed by centrifugation at 11,977 rcf in a Beckman coulter JLA8.1000 rotor at 4 °C. The pellet was washed with 200 ml buffer A (20 mM MES pH 6.5, 10 mM CaCl_2_ 10 mM MgCl_2_) and centrifuged again at 22,000 rcf, 15 min, 4 °C in a JLA16.250 rotor. Pelleted cells were flash-frozen in liquid nitrogen and stored in buffer A supplied with 20% glycerol at -80 °C until further use.

### Purification of Photosystem II and sample preparation

Thylakoid membranes and protein samples were prepared as described previously [24] with some adjustments. After performing the established cell disruption and membrane solubilization methods, proteins were separated by Strep-tag affinity chromatography (AC) and subsequent anion-exchange chromatography (IEC). The previously used Strep-Tactin Superflow hc cartridges were replaced by Strep-Tactin Superflow XT columns (IBA Lifesciences), which makes it possible to replace the Tris pH 7.5 buffer system with MES pH 6.5. This prevents pH changes throughout the whole procedure. PSII was also eluted with biotin (50 mM final concentration) instead of desthiobiotin according to the manufacturer’s instruction.

### Analysis of proteins with polyacrylamide gel electrophoresis

The protein composition of the PSII complexes was analyzed by SDS-PAGE [61] using the Thermo Fisher Scientific Prestained PageRuler #26616 as a size standard. To investigate the oligomeric state of the protein complexes, blue-native PAGE according to Schägger and Jagow [62] using an acrylamide gradient from 3.5 to 16% was done.

### Immunoblot analysis

For immunoblot analysis, proteins were transferred to an Immobilon®-P PVDF membrane (Merck Millipore Ltd.) in a Biometra Fastblot B34 chamber at 1 mA/cm^2^ membrane for 1.5 h in transfer buffer (20 mM CAPSO pH 11.2, 20% (v/v) methanol). Membranes were washed with TBS (10 mM Tris pH 8, 150 mM NaCl) for 10 minutes and blocked in 3% (w/v) BSA in TBS for 2 hours. After washing twice with TBST-T (TBS supplied 0.05% Tween and 0.1% Triton X-100) for 5 min each, the membrane was incubated overnight with streptavidin alkaline phosphatase conjugate (1:5,000; Amersham Biosciences) in TBS supplied with 1% BSA. The membrane was washed twice with TBST and incubated for 10 min in alkaline phosphatase (AP) buffer (100 mM Tris pH 9.5, 100 mM NaCl, 5 mM MgCl_2_). Signals were recorded after incubation with 0.0765 mM nitro blue tetrazolium chloride (NBT) and 0.19 mM 5-bromo-4-chloro-3-indolyl phosphate (BCIP) in AP buffer. The reaction was stopped by washing several times with water.

### Oxygen Evolution experiments

Oxygen evolution rates were determined as described previously [24] using purified PSII with a final chlorophyll concentration of 2-5 µg·ml^-1^.

### Fluorescence measurements

Fluorescence decay and variable fluorescence induction of PSII were measured with an FL3500 Dual-Modulation Kinetic fluorometer (Photon Systems Instruments) as described in [11] and [21], respectively. The samples were prepared in buffer containing 150 mM KCl, 20 mM MES-KOH pH 6.5, 10 mM CaCl_2_, 10 mM MgCl_2_ and 0.03% (w/v) n-dodecyl-ß-d-maltoside (ß-DDM) in the presence or absence of DCMU (3-(3,4-dichlorophenyl)-1,1-dimethylurea or DCBQ (2,6-dichloro-1,4-benzoquinone) at 20 mM with a final concentration of 150 nM PSII reaction centers and measured after 5 min dark adaptation at RT.

The averaged curve of the fluorescence decay data was deconvoluted using the following formula [30],

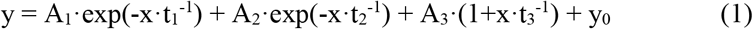

with A_1_, A_2_, A_3_ as the amplitudes with their corresponding time constants t_1_, t_2_ and t_3_. The offset of the y-axis is described by y_0_. For measurements with two instead of three exponents, A_2_ was removed from (1). Half times were calculated by t = ln(2)·t_x_ for the exponential decays, and t = t_x_ for the hyperbolic decay. Since A_1_ is overrepresented during the fluorescence decay, the amplitudes of each phase were determined by their minima and peaks of the respective kinetics in the studied period instead of using the values for A_1_-A_3_.

Kinetic analysis of the variable fluorescence induction data was performed by using the following formula [43] with the same annotations as above for (1). For measurements with two instead of three exponents, A_2_ was removed from (2). Amplitudes were concluded directly from A_1_-A_3_.

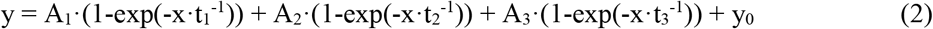

### MALDI-TOF MS

The target subunits were purified and desalted using C18-ZipTips (Merck, Germany) according to the manufacturer’s instructions. The eluted organic fractions were lyophilized in a vacuum concentrator (Eppendorf, Germany), reconstituted in 0.1% Trifluoroacetic acid (TFA) and mixed in a 1:1 ratio with sDHB (Bruker Daltonics, Germany) matrix solution (50 mg·ml^-1^ in 50% Acetonitrile (ACN), 50% water and 0.1% TFA). Subsequently, 1 μl aliquots of the mixture were deposited on a BigAnchor MALDI target (Bruker Daltonics, Germany) and allowed to dry and crystallize at ambient conditions. Unless stated otherwise, all reagents and solvents were obtained from Sigma Aldrich, Germany.

MS spectra were acquired on a rapifleX MALDI-TOF/TOF (Bruker Daltonics, Germany) in the mass range from 3,000-7,000 m/z in reflector positive mode. The Compass 2.0 (Bruker, Germany) software suite was used for spectra acquisition and processing.

### LC-MS

Protein samples were reduced with Tris(2-carboxyethyl)phosphine (TCEP) and cysteines alkylated with Iodoacetamide (IAA, Thermo Fisher, Germany). Subsequent proteolytic digests were performed using S-TRAPs (Protifi, USA) according to the manufacturer’s instructions. Peptides were further desalted and purified on Isolute C18 SPE cartridges (Biotage, Sweden) and dried in an Eppendorf concentrator (Eppendorf, Germany) as described by Zabret et al. [11].

After solubilization in 0.1% formic acid (FA) in ACN/water (95/5, v/v), samples were subjected to LC-MS/MS analysis on an Ultimate 3000 nanoRSLC (Thermo Fisher, Germany) system, equipped with an Acclaim Pepmap C18 trap column (2 cm · 75 µm, particle size: 3 µm; Thermo Fisher) and a C18 analytical column (50 cm · 75 µm, particle size: 1.7 µm; CoAnn Technologies) with an integrated liquid-junction and fused silica emitter coupled to an Orbitrap Fusion Lumos mass spectrometer (Thermo Fisher, Germany). Trapping was performed for 6 min with a flow rate of 6 µl · min^-1^ using loading buffer (98/2 (v/v) water/ACN with 0.05% TFA) and peptides were separated on the analytical column at a flow rate of 250 nl · min^-1^ with the following gradient: 4-48% B in 90 min, 48-90% B in 1 min and constant 90% B for 5 min followed by 18 min column re-equilibration at 4% B with buffer A (0.1% FA in water) and buffer B (0.1% FA in 80/20 (v/v) ACN/water). Peptides eluting from the column were ionized online using a Nano Flex ESI source and analyzed in data-dependent mode. Survey scans were acquired over the mass range from 350-1400 m/z in the Orbitrap (maximum injection time: 50 ms, AGC (automatic gain control) fixed at 2·10E5 with 120K mass resolution) and sequence information was acquired by a top speed method with a fixed cycle time of 2 s for the survey and following MS/MS-scans. MS/MS-scans were acquired for the most abundant precursors with a charge state from 2-10 and an intensity minimum of 3×10E4. Picked precursors were isolated in the quadrupole with a 1.4 m/z isolation window and fragmented using HCD (NCE (normalized collision energy) = 30%). For MS/MS spectra, an AGC of 10E4 and a maximum injection time of 54 ms were used and detection was carried out in the Orbitrap using 30K mass resolution. The dynamic exclusion was set to 30 s with a mass tolerance of 10 ppm. Unless stated otherwise, all reagents and solvents were obtained from Sigma Aldrich, Germany.

Data analysis was performed in Fragpipe 17.1 using MSFragger 3.4 for database searches [63]. Raw files were recalibrated and searched against the Uniprot proteome for *Thermosynechococcus vestitus* BP1 (UP000000440; obtained 2022-03-17) along with the modified PsbO sequence. The search space was restricted to tryptic peptides with a length of 7-50 amino acids, allowing for up to two missed cleavages and with a minimum of one unique peptide per protein group. Carbamidomethylation of Cysteine was set as a fixed modification and oxidation of Methionine as well as N-terminal formylation and acetylation were set as variable modification. Match between runs was enabled and Percolator was used to estimate the number of false-positive identifications. Results filtered for a strict target false discovery rate (FDR) < 0.01.

The mass spectrometry proteomics data have been deposited to the ProteomeXchange Consortium (http://proteomecentral.proteomexchange.org) via the PRIDE partner repository [64] with the dataset identifier PXD033676.

### LC-MS of single bands

To identify the proteins from single SDS-PAGE bands, single bands were extracted, prepared, and digested with trypsin (Promega) as described previously [65]. Samples were measured in a previously described setup [65,66]. In brief, the samples were resuspended in 20 µl solution A (0.1% formic acid (FA), 2% ACN) and applied to a UPLC Symmetry C18 trapping column (5 μm, 180 μm × 20 mm, Waters) afterward transferred to a UPLC BEH C18 column (1.7 μm, 75 μm · 150 mm, Waters) on a nanoAQUITY gradient UPLC (Waters) at a flow rate of 5 µl·min^-1^. The sample was injected into a Thermofisher Scientific Orbitrap ELITE mass spectrometer via a PicoTip Emitter (SilicaTip, 30 µM, New Objective) at a flow rate of 0.4 µl·min^-1^ at 1.5-1.8 kV. The column oven temperature was set to 45 °C. Proteins were eluted over 60 min in a discontinuous gradient of solution A to solution B (0.1% FA in ACN): 0-5 min: 2% B; 5-10 min: 2-5% B; 10-41 min: 5-30 % B; 41-46 min: 30-85 % B; 46-47 min: 95% B; 47-60 min: 2% B. MS1 spectra were recorded at a resolution of 240,000 with a mass range of 300-2,000 m/z in parallel mode by the orbitrap, and MS2 spectra were recorded of the 20 most intense precursors. A dynamic exclusion of 1 min was applied and collision-induced dissociation (CID) was performed at 35% relative energy. Single charged ions were excluded from the analysis.

For analysis, Proteome Discoverer Version 2.3 was used. The spectrum files were recalibrated with a tolerance of 20 ppm and a search was done using Sequest HT with a modified reference proteome database of *Thermosynechococcus vestitus* (UP000000440; obtained 2020-03) [67] with common contaminants from the Global Proteome Machine (GPM) database (https://thegpm.org/crap/index.html; obtained 2020-01) and the modified PsbO-Sequence. Up to two missed cleavage sites and the following modifications were allowed with a peptide length of at least 6 amino acids: N-terminal acetylation, Methionine oxidation. A strict FDR (< 0.01) was applied.

The corresponding mass spectrometry data have been deposited to the ProteomeXchange Consortium (http://proteomecentral.proteomexchange.org) via the PRIDE partner repository [64] with the dataset identifier PXD034220.

### Mn-quantification by EPR

Manganese of isolated PSII was extracted by acid extraction as previously described [29]. In brief, extraction was done by the addition of 0.2 M HNO_3_ and 0.1 M CaCl_2_ to a final concentration of 17.6 µM PSII reaction centers per measurement. After 20 minutes of incubation at 20 °C, the solutions were centrifuged for 15 minutes at maximum speed on a tabletop centrifuge and the supernatants were used for EPR.

All CW EPR measurements were performed on a Bruker X-band (9.86 GHz) E580 (Bruker Biospin) spectrometer equipped with Bruker Super high Q resonator at room temperature. Following measurement parameters were used: power 10 mW, modulation amplitude 10G, modulation frequency 100 kHz, sweep range 1,000 G, sweep time 200 s, number of points 1024. For the measurement, 20 µL of the sample was placed into a 1.5 mM OD glass capillary (Blaubrand), spun down, and sealed with critoseal. All PSII samples were transferred to capillary and measured on the same day after acid extraction. The cavity background signal was measured experimentally on the buffer-only sample and subtracted from the spectra. Mn concentrations of PSII samples were determined by the comparison of double integrals to the calibration curve obtained from MnCl2 stocks at known concentrations. For spectra integration, a home-developed online tool program was used: https://www.spintoolbox.com/. Manganese chloride tetrahydrate MnCl_2_x4H_2_O (Sigma Aldrich) was used to prepare 100 mM stock solution in water and further dilutions were done with extraction buffer.

## Abbreviations

AC: Affinity chromatography
Chl a: Chlorophyll a
EPR: Electron paramagnetic resonance
IEC: Ion exchange chromatography
MALDI-ToF: Matrix-assisted laser desorption/ionization-time of flight
MS: Mass spectrometry
OEC: Oxygen evolving complex
PQ: Plastoquinone
photosystem II: PSII
RC: Reaction center
Twin-Strep-tag: TS-tag

## Statements and Declarations

The authors declare no competing interests.

## Author contributions

M.M.N conceived the study. Generation of the mutant line, protein isolation and biochemical studies were done by J.L. EPR analysis and the corresponding figures were made by S.K. and E.B. Mass spectrometry analysis was done by J.M., J.L. and J.D.L. J.L. and M.M.N. wrote the manuscript and prepared the figures with the contributions of all other authors. All authors approved the final version of the manuscript.

## Acknowledgments and funding

We thank M. Völkel-Lambertz and R. Oworah-Nkruma for excellent technical assistance. This work was funded by the German Research Council (DFG) within the framework of the research unit FOR2092 (NO 836/3-2 to M.M.N.) and the DFG priority program 2002 (NO836/4-1 to M.M.N. and 3542/1-1 to J.D.L.).

