## Supplemental Information for "Isolation of a novel heterodimeric PSII complex via strep-tagged PsbO"

### Supporting Information

#### Additional files

**SI File 1: MS Table 1**: Single band LC-MS/MS analysis of the additional protein band found in saD.

**SI File 2: MS Table 2**: LC-MS/MS analysis of the whole protein complexes.

#### Supplementary figures and tables

**SI Table 1: MALDI-TOF analysis overview** in the range of 3,000-10,000 m/z. Assigned proteins and their theoretical values and masses assigned in previous studies [1,2]. The measured masses are shown on the right. Assigned modifications are shown in parentheses.

| Name | Theoretical average masses (Uniprot full seq; published) | Measured monoisotopic masses (rounded) / assigned mass (+mods) |
|---|---|---|
| PsbT | 3,875; 3,906/3,904 (n-formyl) | 3,900 (formyl); 3,916 (formyl + Ox) |
| PsbJ | 4,105; 4,002 (-M + n-formyl) | 3,999 (-M); 4,013 (-M + Ox) |
| PsbM | 3,981; 4,011/4,009 (n-formyl) | 4,006 (formyl); 4,022 (formyl + Ox) |
| PsbK | 5,026; 4,103/4,101 (1-9 propeptide removed) | 4,097 |
| PsbX | 4,319; 4,192/4,190 (-M) | 4,186 |
| PsbL | 4,297; 4,301/4,299 | 4,294; 4,310 (Ox) |
| PsbI | 4,405; 4,437/4,435 (n-formyl) | 4,431 (formyl); 4,447 (formyl + Ox) |
| PsbE | 9,573; 9,446/9,440 (-M) | 4,718 (2+ charged) |
| PsbF | 5,065; 4,981/4,977 (-M +n-acetyl) | 4,973 (-M +n-acetyl); 4,989 ((-M +n-acetyl)+Ox) |
| PsbU | 15,018; 11,649/11,641 (1-30 signal peptide removed) | 5,820 (2+ charged) |
| PsbZ | 6,764; 6,793/6,798 (n-formyl) | 6,789 (formyl) |

**SI Table 2: Table of spectral areas of 1st and 2nd extractions** of fully assembled PSII aD samples obtained after double integration of background-corrected spectra (Figure SI 5). Corresponding manganese concentrations were calculated by the division to the slope of calibration curve (Figure SI 4c). SD represents the sample standard deviation obtained from different biological (1-4) and technical (X-2) replicates. Extraction 2 represents an additional extraction step with the pellet of the first extraction to confirm the efficiency of the extraction.

| | | Extraction 1 | | Extraction 2 | | Total |
|---|---|---|---|---|---|---|
| | Sample | Spectral area | Concentration (μM) | Spectral area | Concentration (μM) | |
| 1 | aD 1 | 5,347 | 66.79 | 223 | 2.79 | |
| 2 | aD 2 | 5,136 | 64.15 | 168 | 2.10 | |
| 3 | aD 3 | 4,948 | 61.80 | 149 | 1.86 | |
| 4 | aD 4 | 4,794 | 59.88 | 143 | 1.79 | |
| 5 | aD 1-2 | 5,609 | 70.06 | 235 | 2.94 | |
| 6 | aD 2-2 | 5,662 | 70.72 | 240 | 3.00 | |
| 7 | aD 3-2 | 5,310 | 66.33 | 251 | 3.14 | |
| 8 | aD 4-2 | 5,170 | 64.58 | 203 | 2.54 | |
| | mean | | 65.54 | | 2.52 | 68.06 |
| | SD | | 3.75 | | 0.53 | 3.79 |
| | Mn/cluster | | 3.72 | | 0.14 | 3.87 |
| | SD | | 0.21 | | 0.03 | 0.21 |

**SI Table 3: Table of spectral areas of 1st extraction of saD and aM** obtained after double integration of the spectra after background subtraction (Figure SI 6). Corresponding manganese concentrations were calculated by the division to the slope of calibration curve (Figure SI 4c). SD represents the sample standard deviation obtained from different biological (1-5) and technical (X-2) replicates.

| | Sample | Spectral area | Concentration (µM) | Sample | Spectral area | Concentration (µM) |
|---|---|---|---|---|---|---|
| 1 | saD 1 | 4,269 | 53.32 | aM 1 | 4,875 | 60.89 |
| 2 | saD 2 | 3,749 | 46.83 | aM 2 | 4,691 | 58.59 |
| 3 | saD 3 | 3,739 | 46.70 | aM 3 | 4,685 | 58.52 |
| 4 | saD 4 | 3,241 | 40.48 | aM 4 | 4,471 | 55.85 |
| 5 | saD 1-2 | 4,316 | 53.91 | aM 1-2 | 4,963 | 61.99 |
| 6 | saD 2-2 | 3,979 | 49.70 | aM 2-2 | 4,527 | 56.55 |
| 7 | saD 3-2 | 3,798 | 47.44 | aM 3-2 | 4,784 | 59.76 |
| 8 | saD 4-2 | 3,800 | 47.46 | aM 4-2 | 5,110 | 63.83 |
| 9 | saD 5 | 3,379 | 42.21 | | | |
| 10 | saD 5-2 | 3,591 | 44.85 | | | |
| | mean | | 47.29 | | | 59.50 |
| | SD | | 4.28 | | | 2.69 |
| | Mn/cluster | | 2.69 | | | 3.38 |
| | SD | | 0.24 | | | 0.15 |

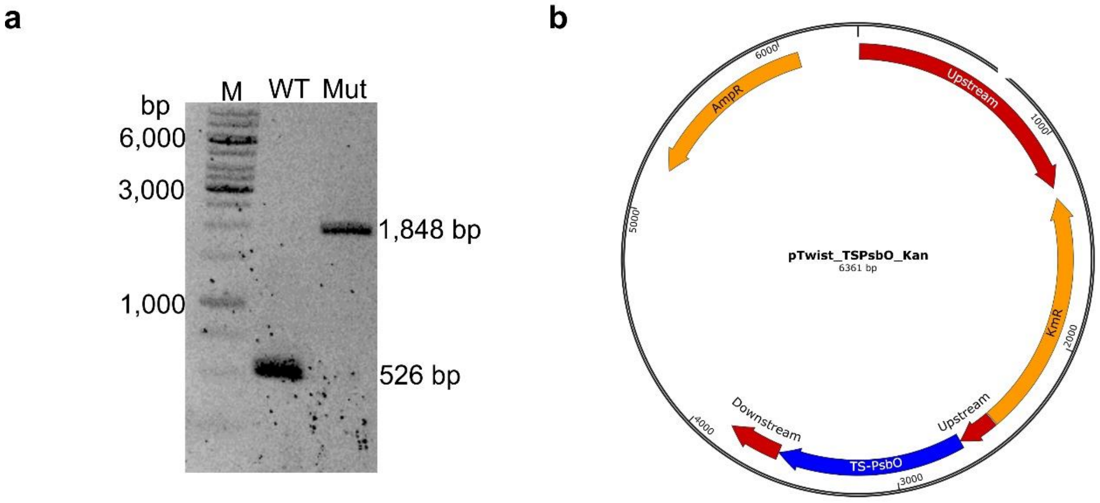

**Figure SI 1: Cloning information.**

**a)** Segregation check of the TS-PsbO mutant (Mut) compared to the wild type strain (WT). Primers were designed to bind in the upstream region in front of the inserted the Kanamycin-resistance cassette and in the PsbO-gene. M: size standard Thermo Scientific GeneRuler 1 kb. **b)** Plasmid used for transformation. The plasmid was designed with Clone Manager and synthesized by Twist Bioscience in their pTwist vector. Graphic visualized with Snap Gene Viewer.

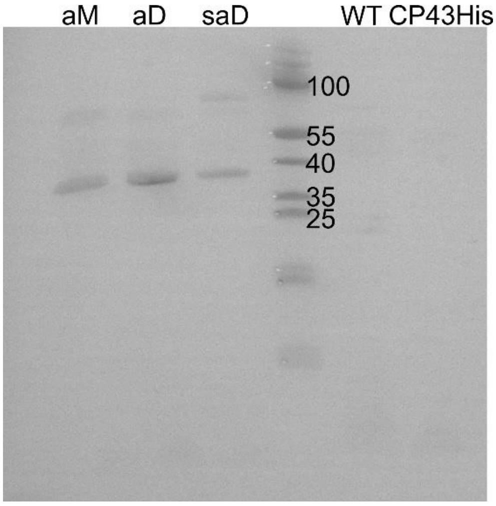

**Figure SI 3: Western Blot against the Twin-Strep-tag linked to the PsbO subunit.**

Per lane, PSII with a Chl-content of 2.5 µg were applied. Controls: active PSII dimer from wild type [3] or CP43His-Tag preparations [4]. As size standard, Thermo Fisher Scientific PageRuler #26616 was applied.

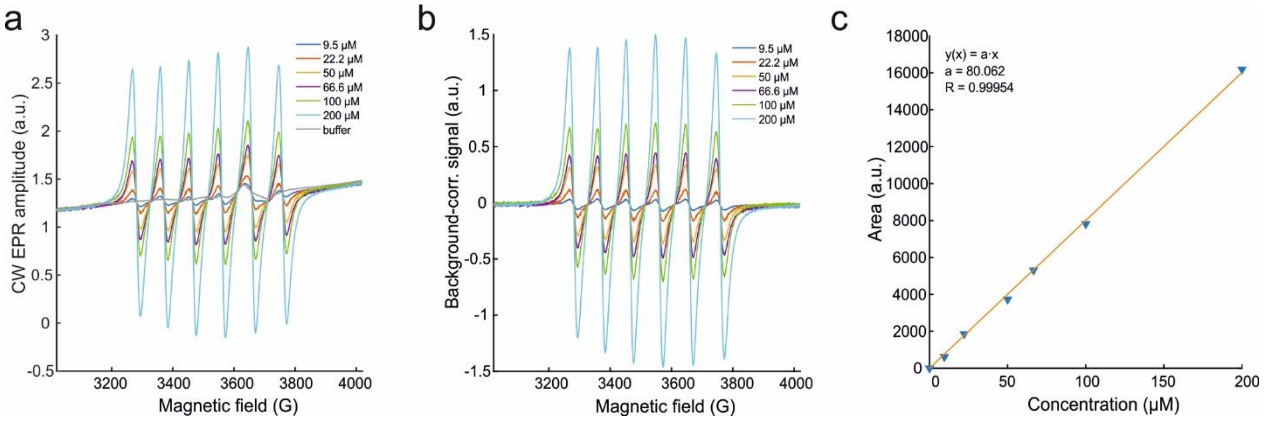

**Figure SI 4**: **X-band (9.86 GHz) continuous wave (CW) EPR spectra of $MnCl_2$ stock solutions at different concentrations.**

**a)** Raw and **b**) background-corrected spectra. For the background correction, the spectrum of the buffer (grey in **a**) was subtracted. **c)** Calibration curve obtained as double integrals of spectra shown in **b** plotted against $MnCl_2$ concentrations. The corresponding fit a·x in **c** shows a linear dependence on concentration with the slope a = 80.06 $\mu M^{-1}$. The unknown Mn-concentrations of aD, aM, and saD samples were calculated as corresponding double integral values divided by the slope.

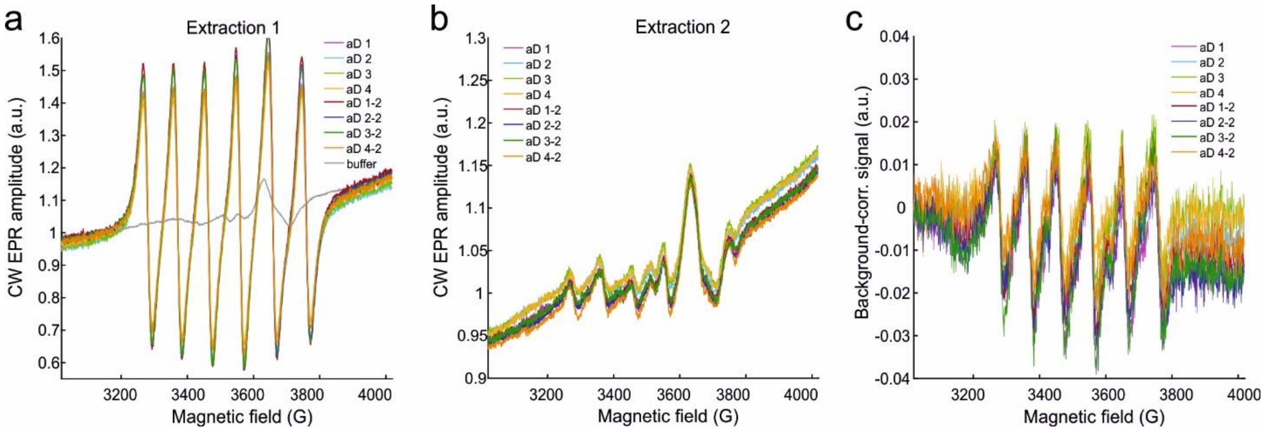

**Figure SI 5**: **X-band (9.86 GHz) continuous (CW) EPR spectra (left panels) of aD**

RC concentration: $C_{RC}$ =17.6 µM. Raw data after the first (**a**) and second (**b**) acid extraction. The second reaction was done with the pellet of the first extraction to confirm the efficiency of the extraction (see also SI table 2). The buffer spectrum (grey in **a**) was used for subsequent background subtraction (**c**).

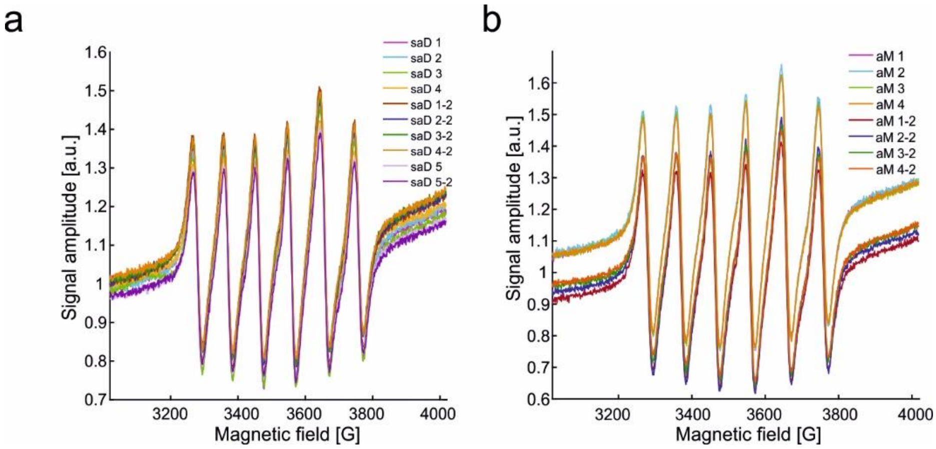

**Figure SI 6: X-band (9.86 GHz) continuous wave (CW) EPR spectra** of saD **(a)** and aM **(b)** shown as detected. RC concentration: $C_{RC}$ =17.6 µM.

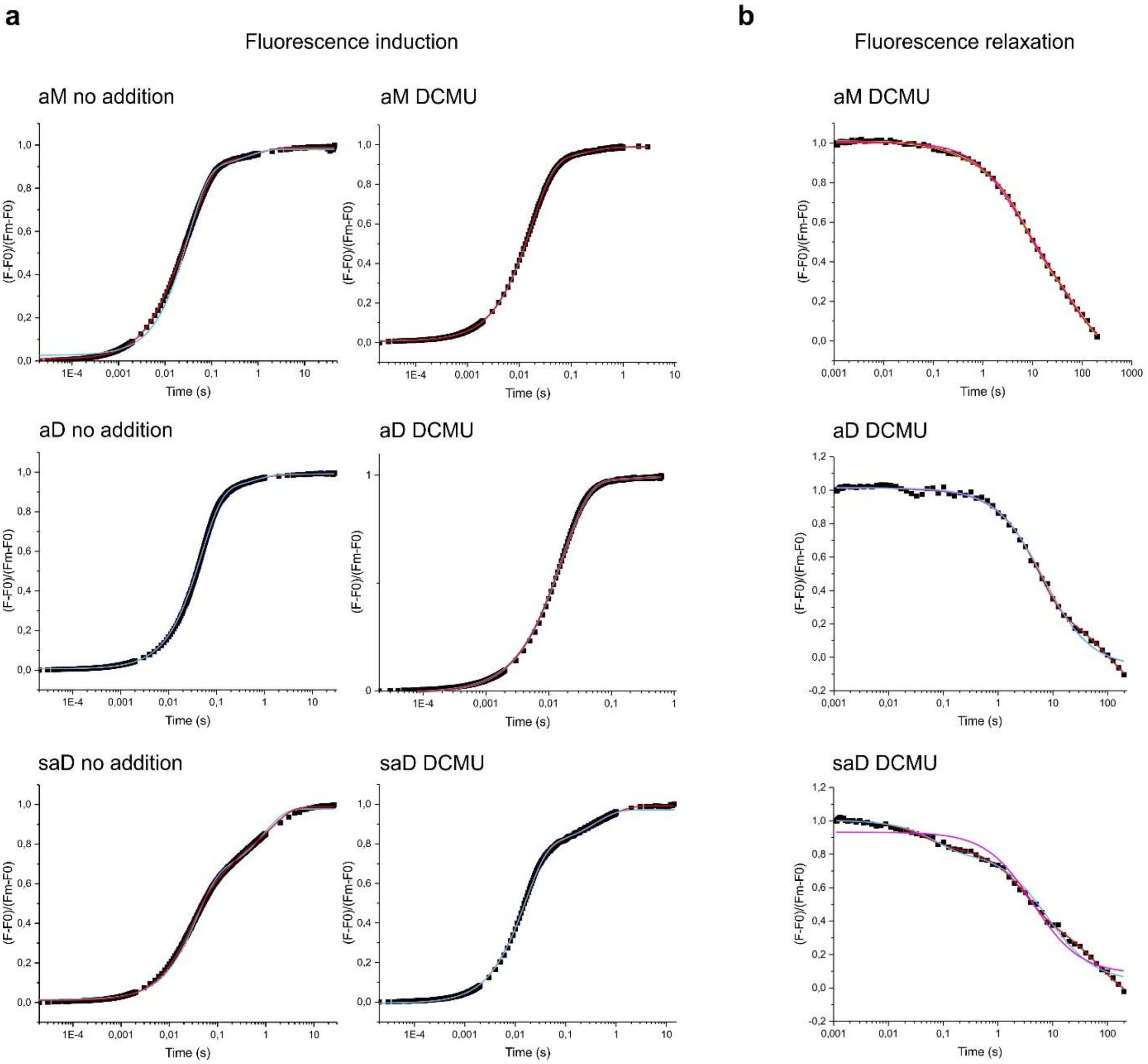

**Figure SI 7: Selected fits of the Chl a fluorescence kinetic measurements**

**a)** Fluorescence Induction measurements of aM, aD and saD in absence and presence of DCMU and fitting by using two (red) or three exponents (light blue). Averaged data is shown in black dots. **b)** Fluorescence decay kinetics of the three complexes in presence of DCMU. Red: two exponential and one hyperbolic decay. Light blue: one exponential and one hyperbolic decay. Pink: two hyperbolic decays. Yellow: one exponential and two hyperbolic decays.
